# Industrialization is associated with elevated rates of horizontal gene transfer in the human microbiome

**DOI:** 10.1101/2020.01.28.922104

**Authors:** Mathieu Groussin, Mathilde Poyet, Ainara Sistiaga, Sean M. Kearney, Katya Moniz, Mary Noel, Jeff Hooker, Sean M. Gibbons, Laure Segurel, Alain Froment, Rihlat Said Mohamed, Alain Fezeu, Vanessa A. Juimo, Catherine Girard, Le Thanh Tu Nguyen, B. Jesse Shapiro, Jenni M. S. Lehtimäki, Lasse Ruokolainen, Pinja P. Kettunen, Tommi Vatanen, Shani Sigwazi, Audax Mabulla, Manuel Domínguez-Rodrigo, Roger E. Summons, Ramnik J. Xavier, Eric J. Alm

## Abstract

Horizontal Gene Transfer, the process by which bacteria acquire new genes and functions from non-parental sources, is common in the human microbiome ^1,2^. If the timescale of HGT is rapid compared to the timescale of human colonization, then it could have the effect of ‘personalizing’ bacterial genomes by providing incoming strains with the genes necessary to adapt to the diet or lifestyle of a new host. The extent to which HGT occurs on the timescale of human colonization, however, remains unclear. Here, we analyzed 6,188 newly isolated and sequenced gut bacteria from 34 individuals in 9 human populations, and show that HGT is more common among bacteria isolated from the same human host, indicating that the timescale of transfer is short compared to the timescale of human colonization. Comparing across 9 human populations reveals that high rates of transfer may be a recent development in human history linked to industrialization and urbanization. In addition, we find that the genes involved in transfer reflect the lifestyle of the human hosts, with elevated transfer of carbohydrate metabolism genes in hunter gatherer populations, and transfer of antibiotic resistance genes among pastoralists who live in close contact with livestock. These results suggest that host-associated bacterial genomes are not static within individuals, but continuously acquire new functionality based on host diet and lifestyle.

## Main text

Gut bacteria living in symbiosis with humans have experienced high rates of horizontal gene transfer (HGT) over evolutionary time, at least across individuals in industrialized countries ^1,2^. Yet it remains unclear how rates of HGT compare to typical bacterial residence time in the human gut, and how the lifestyle of the human host might influence the rate of HGT and the type of genes transferred.

If the timescale of transfer is slower than within-host residence time, then individual microbiomes will primarily acquire new functions through the acquisition of new strains. However, if the rate of transfer is sufficiently rapid, then a microbiome that is ‘stable’ in terms of bacterial populations ^3–5^ could nonetheless evolve in response to host-specific environmental perturbations through HGT, perhaps in response to diet or changes in cultural practices.

Specific examples demonstrate that HGT can occur within a single individual ^6–10^, especially when there is strong selection for target functions such as antibiotic resistance ^11–13^. But what fraction of species in the human microbiome have acquired genes from another species in their most recent human host, and how does the timescale of HGT compare to the timescale of human colonization? In our previous study ^1^, we focused on HGTs involving sequences with similarity higher than 99% and length higher than 500bp. By assuming a typical molecular clock of ~1 SNP/genome/year ^14^ and genome size of 10^6^ bp, these criteria are consistent with transfer events that happened between 0 and 10,000 years ago. Thus, to answer the question of whether commensal strains routinely acquire new functionality through HGT, more precise estimates of the timescale for HGT are needed.

To measure the rate of HGT on shorter timescales, we compared the amount of transfer observed between bacteria isolated from within the same individual with that observed between bacteria from different individuals. We hypothesized that if the rate of transfer was fast compared to the typical residence time of a bacterial lineage colonizing the human body, then we would observe higher levels of transfer between strains isolated from the same host. Alternatively, if the timescale for transfer was sufficiently longer than a human lifespan, then we would observe similar levels between bacteria regardless of whether they were isolated from the same host. To focus our analysis on the most recent events, we looked for large blocks (>10kb) of 100% identical DNA, corresponding to HGT events that occurred between 0 and ~100 years ago, though we also confirm our findings using shorter mobile elements with length larger than 500bp. In this study, we focus only on transfers occurring between bacterial species, ignoring within-species gene recombination events.

Existing reference isolate genomes ^4,15–19^ cannot be used to test for direct gene transfer between any two bacteria within people, because nearly all of those strains were isolated from different individuals. In addition, these reference collections were sampled almost exclusively from industrialized populations, and do not reflect the diversity of human lifestyles. Therefore, we analyzed the whole genomes of 6,188 newly cultured bacterial isolates using stool samples collected from 34 individuals in 9 human populations worldwide: the Hadza and Datoga in Tanzania, Beti and Baka populations in Cameroon, Inuit individuals in Canadian Arctic, Sami and Finnish individuals in Finland, and individuals from a Northern Plains Tribe in Montana and from the Boston area in the USA; Supplementary Figure 1 & Supplementary Table 1 for a description of lifestyles). We grouped bacterial genomes into species clusters based on genomic similarity (using the Mash distance as a proxy of the Average Nucleotide Identity, see Methods). These genomes represent 253 bacterial species across 6 phyla, grouping into 62 known and 54 unknown genera (Figure 1A & Supplementary Tables 2 & 3 for culturing data and genome assembly statistics). The sampled human populations had different genetic backgrounds and very different lifestyles, ranging from industrialized to hunter-gatherer communities. We sampled many bacterial isolates of different species within each individual, and detected thousands of recent HGTs in our genomic data: in total, we captured 134,958 mobile elements across multiple bacterial species, both within and between people. 57% of the bacterial genomes (3556/6188) were involved in at least one recent HGT event (Figure 1A), indicating that HGT is rampant in the contemporary human gut.

**Figure 1.**
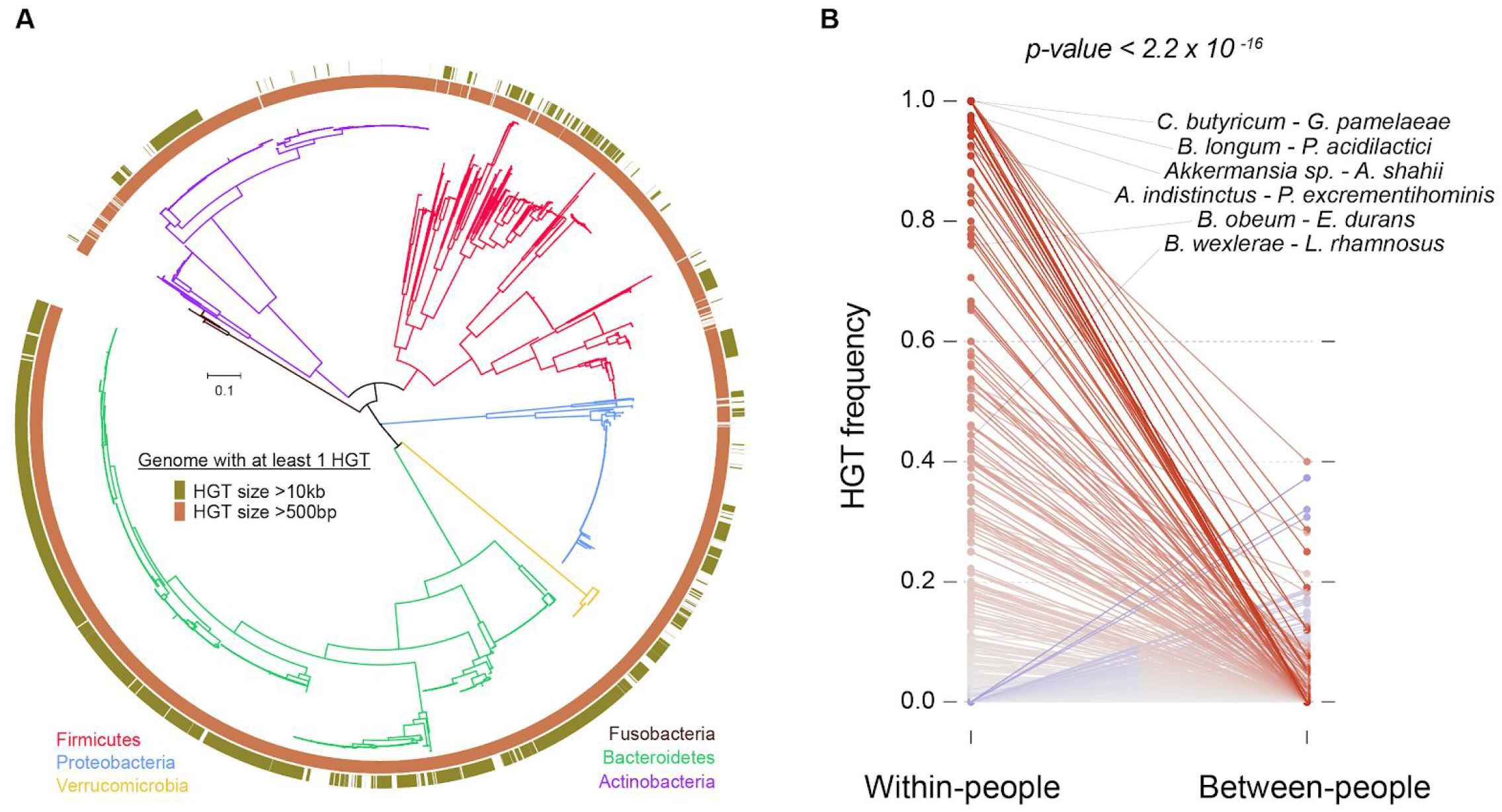
HGT is common within the gut microbiome of individual people. **(A)** Phylogenomic tree of the 6,188 human gut bacterial isolates that we generated in this study and that were sampled from 9 human populations. Branches are colored by phylum. The inner and outer rings show genomes in which at least 1 HGT larger than 500bp and 10kb was detected, respectively. **(B)** HGT frequencies within and between people were computed using the whole set of genomes. Solid lines represent bacterial species pairs sampled both within and between individuals. Differences in HGT frequency are colored along a gradient from grey (no difference) to red (within-people HGT frequency is higher than between-people) or from grey to blue (between-people HGT frequency is higher than within-people), darker colors representing higher differences. The HGT frequency of bacterial species pairs found within people was compared to the expected frequency based on the HGT frequency of the same species pairs found in different people (*p-value* < 2.2×10^−16^). Observed and expected HGT frequencies were calculated using the total number of genome comparisons with at least 1 HGT (see Methods). A few distantly-related species pairs that exchange genes within people at higher frequency than we could expect by phylogeny (see Figure 2A) are listed.

We found that bacterial species pairs sampled within people are more likely to share recently transferred DNA than the same species pairs sampled from two different persons (the number of observed within-person HGT events was compared to the expected number of events based on the number of between-person events, correcting for species composition and uneven sampling depth, Figure 1B, *p-value* < 2.2×10^−16^, see Methods), and this signal is driven by many different bacterial species covering diverse taxonomic groups (Figure 1A & 1B). This result suggests that the timescale for HGT is short. Strictly speaking, we cannot distinguish between transfers that occurred in the host of origin from those that may have occurred in a host’s parent or even grandparent. However, it is unlikely that a large fraction of transfers occurred prior to colonization of the host because the overall rate of HGT is large compared to the rate of inheritance of strains from a parent (see discussion in Supplemental Information). These results are robust to the particulars of our analysis: an increase in HGT frequency within individuals is replicated when restricting analyses to within each of our sampled populations, or when considering the 5,126,962 mobile elements larger than 500bp that are distributed across 98% (6068/6188) of our genomes (*p-value* < 2.2×10^−16^) (Figure 1A & Supp Fig. 2 & 3). Together, these results suggest that HGTs occur on timescales that are sufficiently short to reshape gut community functions extensively and continuously during an individual’s lifetime.

Because HGT frequency is primarily driven by transfers occurring among closely-related organisms, which tend to exchange more genes together than distantly-related species, we investigated HGT frequency over a range of phylogenetic distances. We show that phylogenetic relatedness is a strong driver of HGTs overall (more closely related species transferring more genes, Linear Mixed Effects model fit test, *p-value* < 2.2×10^−16^), and that the strong enrichment for transfer within individuals as compared to between individuals occurs across all phylogenetic distances (Figure 2A), which holds true even when considering all HGTs larger than 500bp (Supplementary Figure 4).

**Figure 2.**
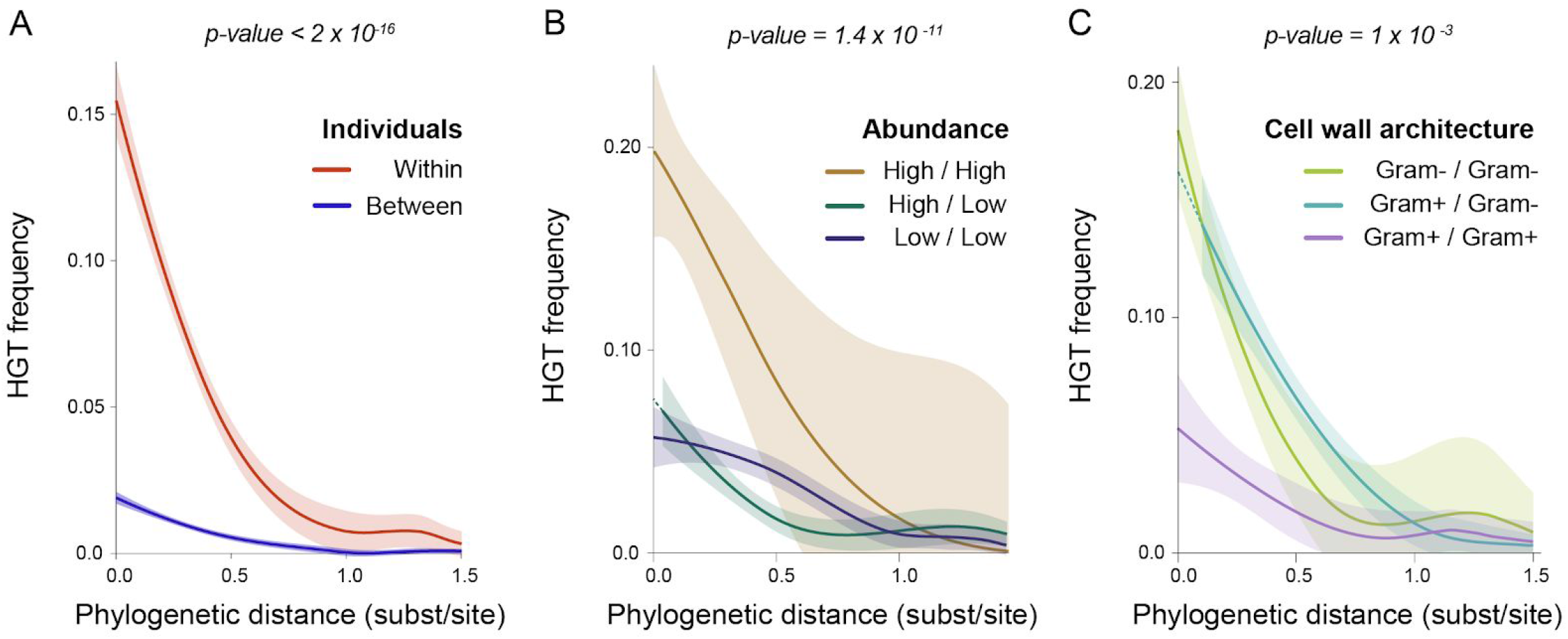
Phylogeny, abundance and cell wall architecture drive gene transfer. The individual contributions of phylogeny, abundance and cell wall architecture were measured using a Linear Mixed Effects model and plotted using loess regressions, with confidence intervals being calculated from the standard errors. P-values associated to each factor are shown above each plot. (**A**) HGT frequency within people is higher than between people across all phylogenetic distance bins. Phylogenetic distances were derived from the phylogenomic tree in Figure 1A. A few distantly-related species pairs that exchange genes within people at higher frequency than we could expect by phylogeny are highlighted in Figure 1B. (**B**) HGT frequency is plotted across species abundance bins. Bacterial abundances are individual-specific, and were measured by mapping metagenomic reads against individual genomes (see Methods). We used a threshold of 0.01 to define highly and lowly abundant bacteria. The HGT frequency is linearly extrapolated for the High/Low category in the range of very small phylogenetic distances (dashed line) due to the absence of species pairs with closely-related species in this category. (**C**) HGT frequency is plotted across types of cell wall architecture. We used Gram staining as a proxy to call for monoderm or diderm bacteria. As in B, the dashed line extrapolates the HGT frequency for the Gram+/Gram− category, as no species pairs with small phylogenetic distances were sampled within this category.

Having established the rapid timescale of HGT, we next asked what factors drive gene exchange frequency in the human gut. We hypothesized that pairs of highly abundant species in a given ecosystem would have a higher probability of gene exchange compared to pairs involving at least one low-abundance species, independent of their phylogenetic distance, though we previously argued against a major role for abundance in controlling HGT frequency ^1^. This hypothesis had never been directly tested because datasets that paired in-depth genomic sampling with accurate abundance estimates did not yet exist. To test the abundance hypothesis, we generated metagenomic data for the stool samples from which we had cultured bacterial isolates, and calculated the average abundance of each bacterial species within each person by mapping metagenomic reads against the isolate genomes (see Methods). We found that species abundance is a strong determinant of HGT (Linear Mixed Effects model fit test, *p-value* = 1.4×10^−11^), independent of phylogeny (Figure 2B), which is replicated when looking at all HGTs larger than 500bp (Supplementary Figure 5). Abundant bacteria are more likely to engage in HGT with other abundant bacteria, which is consistent with the canonical mechanisms of HGT (e.g. conjugation, transformation and transduction ^20^) that involve cell-to-cell contact or access to free DNA in the environment.

Since HGT is driven by phylogenetic distance and abundance, and abundance is similar across individuals within a host population ^5^, we hypothesized that the same gut bacterial species would exchange genes across individuals. To test this hypothesis, we compared HGT frequencies for bacterial species pairs shared by a minimum of 4 individuals within our USA cohort. We found that HGT frequency is homogeneous across people for the majority of bacterial species (the observed average standard deviation of within-person HGT frequency across people was compared to the expected distribution using a randomization test with 1,000 permutations, *p-value* < 0.001, Supplementary Figure 6). This suggests that the core set of abundant lineages shared by individuals within a given population represents a core network of gene exchange that allows bacterial lineages to adapt to common selective pressures acting in the host population.

We next asked whether the architecture of cell envelopes contributes to differences in HGT frequency, independent of phylogeny and abundance. We used reference Gram staining data for each bacterial species as a proxy of cell wall architecture, in order to separate gram-positive monoderm bacteria (single cytoplasmic membrane and a thick peptidoglycan layer) from gram-negative diderm bacteria (two membranes surrounding a thin peptidoglycan layer). We found that diderm bacteria engage more frequently in HGTs than monoderm bacteria, independently of phylogeny and abundance (*p-value* = 1×10^−3^, Figure 2C), which is also observed when considering all HGTs larger than 500bp (Supplementary Figure 7). Interestingly, HGT frequency between two diderm bacteria was similar to HGT frequency between a monoderm and a diderm bacteria, suggesting that diderm bacteria have transfer mechanisms that allow them to share DNA material with a much broader spectrum of genetic backgrounds.

Transitioning from non-industrialized to industrialized lifestyles is associated with drastic changes in microbiome diversity and composition ^21–23^. However, little is known about how these lifestyle transitions impacted the patterns of gene exchange in the human gut microbiome.

To test whether human populations with an industrialized lifestyle have different HGT patterns when compared to populations with non-industrialized lifestyles, we looked at the species pairs in our dataset that are shared by individuals living in the USA (Boston area) and individuals living in either one of the four populations from which we have the largest sampling of bacterial species: the Hadza (hunter-gatherers), the Datoga (pastoralists), the Beti (agriculturalists) and the Baka (currently transitioning from a hunter-gatherer to an agriculturalist lifestyle). For each bacterial species pair, we computed the average HGT frequency at the human population level, looking at shared identical (100%) DNA blocks that are larger than 500bp. Surprisingly, we found that species pairs sampled in the US industrialized population exchanged genes more frequently than when they are found in non-industrialized populations (the number of observed non-industrialized population HGT events was compared to the expected number of events based on the number of industrialized population events, correcting for species composition and uneven sampling depth, *p-value* < 2.2×10^−16^, see Methods) (Figure 3A). This effect holds when restricting the analysis to each non-industrialized population individually compared to the US (Figure 3B). Taken together, these results show for the first time that host lifestyle shapes gene transfer frequencies in the human gut microbiome. These results also suggest that transitioning to industrialized lifestyles resulted in a drastic increase in gene transfers within the gut microbiome, potentially due to increased environmental perturbations to gut bacterial populations.

**Figure 3.**
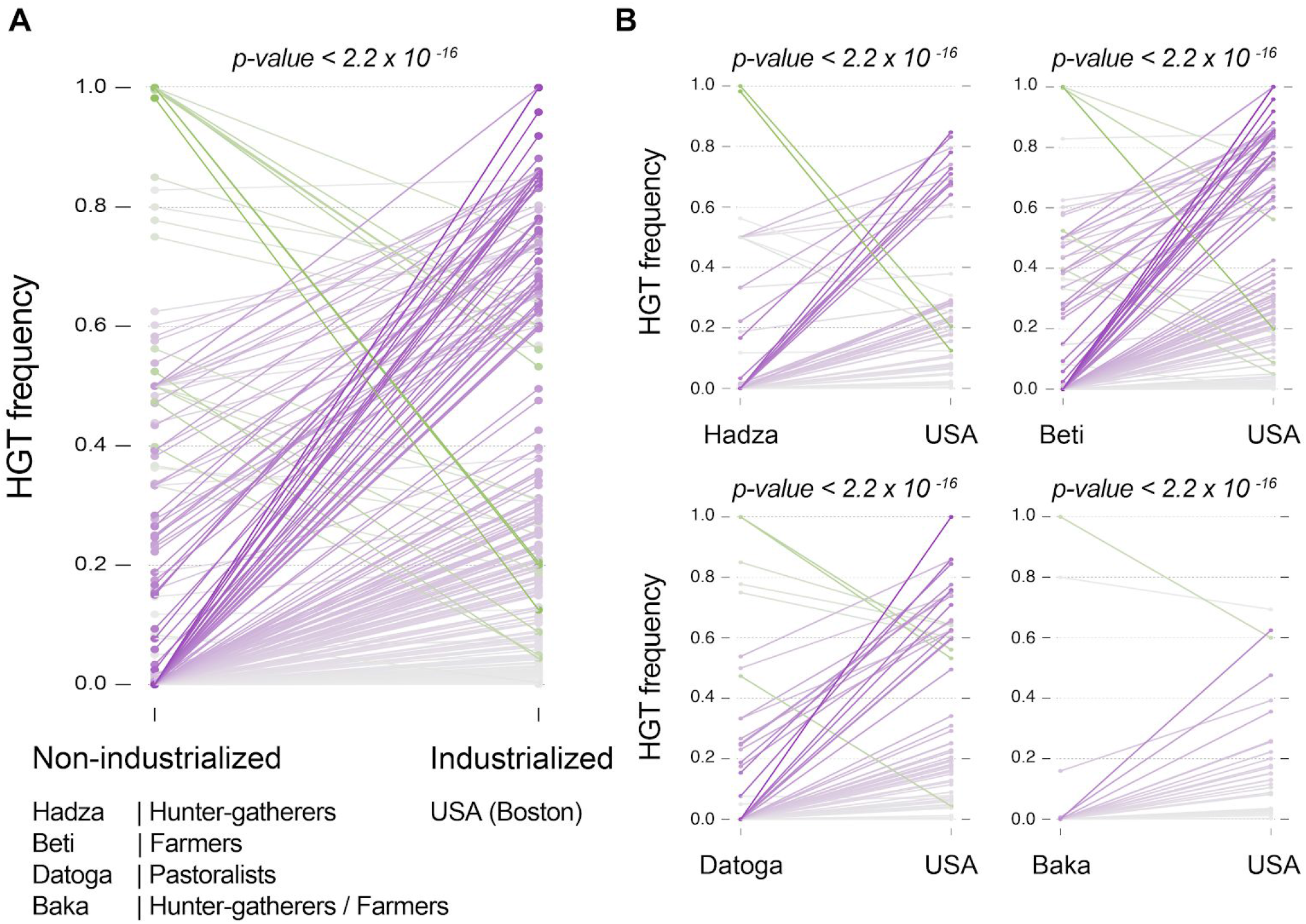
Higher HGT frequency in the gut microbiome of individuals living in industrialized populations. We compared the HGT frequency of all species pairs shared between the USA cohort (industrialized people) and four non-industrialized African cohorts (Hadza people, hunter-gatherers; Beti people, farmers; Datoga people, pastoralists; and Baka people, hunter-gatherers and farmers). (**A**) Comparison of HGT frequencies between the USA cohort and the four aggregated non-industrialized cohorts. Each line represents a species pair found in both the industrialized and non-industrialized groups. Differences are colored along a gradient from grey (no difference) to purple (HGT frequency is higher in USA individuals) or from grey to green (HGT frequency is higher in non-industrialized individuals), darker colors representing higher differences. The number of observed non-industrialized population HGT events was compared to the expected number of events based on the number of industrialized population events (*p-value* < 2.2 x 10^−16^), correcting for species composition and uneven sampling depth. Importantly, results are replicated when species pairs having higher abundance in the USA are removed from the analysis (p-value < 2.2 x 10^−16^), to control for the effect of abundance on HGT frequency. (**B**) Gut bacterial species in USA individuals exchange genes at higher frequency than in non-industrialized communities, consistently across the four non-industrialized ethnic groups (all *p-values* < 2.2×10^−16^).

We reasoned that if HGT occurs on very short timescales, then the type of genes being transferred should reflect the unique selective pressures associated with different individual hosts and populations ^24^. Using gene transfers involving species pairs found in both the USA population and either the Hadza, Beti or Datoga peoples, we first compared broad functional category profiles and found that they differed across lifestyles (Figure 4A, chi-square Goodness-of-fit test, *p-values* < 0.001).

**Figure 4.**
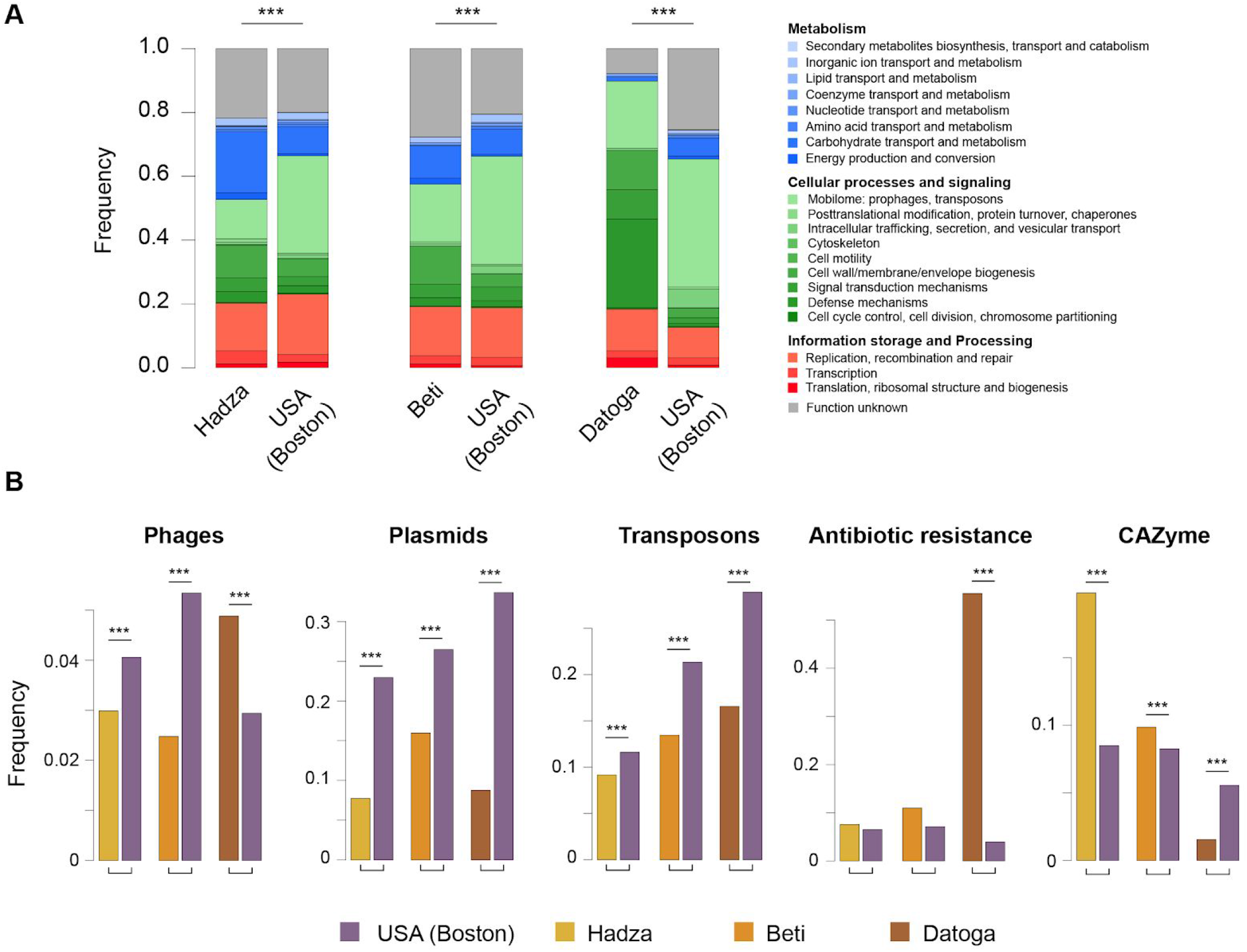
Strong association between host lifestyle and transferred gene functions. Genes within mobile elements were annotated using a variety of reference gene function databases (see Methods) to compare functional profiles of transferred genes between industrialized and non-industrialized populations. Only host populations with a sufficient number of genes annotated with known predicted functions were included in the analysis (USA, Hadza, Beti and Datoga communities; Baka individuals were removed). To account for differences in species composition, HGT functions were counted using only species pairs that are shared by the two compared host populations (USA vs. a non-industrialized population) being compared. For this reason, functional profiles for USA slightly change across pairwise population comparisons. (**A**) Profiles of COG functional categories were compared using a chi-square Goodness-of-fit test (***: *p-values* < 0.001). (**B**) HGT counts of phage, plasmid, transposon, antibiotic resistance and CAZyme genes were compared between industrialized and non-industrialized host populations using two-proportions Z-tests and a Bonferroni correction for multiple tests (***: *p-values* < 0.001).

Having shown that broad functional differences exist across the types of genes transferred in different populations, we focused on genes involved in functions that we thought may differ across populations, including genes involved in mobile elements (phage, plasmid, transposon), antibiotic resistance and carbohydrate-degrading (CAZyme) functions. We found that gut bacteria in industrialized populations exchanged higher relative amounts of plasmid, transposon and phage elements (Figure 4B, two-proportions Z-tests, corrected *p-values* < 0.001), consistent with overall higher levels of HGT. Hadza and Beti individuals, who consume large amounts of non-digestible fibers, host gut bacteria that exchange CAZyme genes at higher frequencies than individuals living in the USA (Figure 4B). Very high transfer frequencies of antibiotic resistance genes were also found in the gut microbiomes of Datoga individuals. The Datoga are pastoralists, raising primarily cattle, and consuming high levels of meat and dairy products from their animals. Like other pastoral farmers in northern Tanzania, they administer antibiotics to their herds ^25,26^. Our results suggest that these recent agricultural practices rapidly altered the fitness landscape in the guts of Datoga people and have already impacted the patterns of gene transfers within their microbiomes. As the use of commercial antimicrobials is now widespread among pastoralist populations in developing countries, similar effects may occur in many populations worldwide with broader impact on the spread of antimicrobial resistance outside the clinic.

Numerous studies have investigated how changes in diet and clinical interventions such as fecal microbiota transplants ^27,28^ impact the composition of the gut microbiome. But inferring mechanistic understanding from compositional changes is difficult. Our study reveals that HGTs within the gut microbiome reflect the unique selective pressures of each human host. Thus, HGT patterns can then be used to identify selective forces acting within each individual and to gain a more mechanistic understanding of these events. Our results also show that whole genome sequencing data provides information on personalized microbiome function at a level of precision that popular approaches, such as 16S amplicon and metagenomic sequencing, cannot achieve. Finally, the high rate of HGT in the human gut is likely a recent development in response to industrialized lifestyle, which was further accompanied by drastic changes in the nature of genes being exchanged. We may not yet fully appreciate the consequences of these shifts in HGT frequency and function on human health.

## Acknowledgements

We are grateful to our field collaborators in Montana (US), Canada, Finland, Cameroon and Tanzania. We thank all human communities that agreed to provide samples to the Global Microbiome Conservancy project. This work was supported by grants from the Center for Microbiome Informatics and Therapeutics at MIT and the Rasmussen Family Foundation, and by a BroadNext10 award from the Broad Institute. Additional support was provided by a Marie Skłodowska-Curie fellowship (A.S. - H2020-MSCA-IF-2016-780860). We thank Tamara Mason and the team at the Walkup Sequencing platform at the Broad Institute for support on sequencing efforts.

## Author contributions

M.G., M.P. and E.J.A. designed this study. M.G., M.P., A.S., K.M., R.E.S., R.J.X. and E.J.A founded the Global Microbiome Conservancy project under which field collections occurred. M.G., M.P., A.S., M.N., J.H., S.M.G., L.S., A.F., R.S.M., A.F., V.A.J., C.G., L.T.T.N., B.J.S., J.M.S.L., L.R., P.P.K., T.V., S.S., A.M. and M.D-R. managed field administrative work and performed the collection of data and samples. M.G. performed computational work and data analyses. M.P. performed the culturing, DNA extraction and sequencing work. S.M.K. performed gene annotations on transferred mobile elements. M.G., M.P. and E.J.A. analyzed the results. M.G., M.P. and E.J.A. wrote the manuscript, which was improved by all authors.

## Methods

### Data availability

Data will be made available online upon acceptance of this manuscript

### Study cohorts, sample collection and storage

Stool samples from 11 North American individuals from the Boston Area (Massachusetts) were obtained from OpenBiome (https://www.openbiome.org/), a non-profit stool bank, under a protocol approved by the Institutional Review Boards (IRB) at MIT and the Broad Institute (IRB protocol #1603506899). Subjects were healthy Fecal Microbiota Transplantation donors screened by OpenBiome to minimize the risk for carrying pathogens. Raw stool was diluted 1:10 in 12.5% glycerol buffer and 0.9% NaCl, homogenized and filtered through a 330um filter.

Stool samples from 23 individuals recruited worldwide as part of the Global Microbiome Conservancy project (microbiomeconservancy.org) were obtained from Inuit people in Canadian Arctic, Sami and Finnish peoples in Finland, Beti and Baka peoples in Cameroon, Hadza and Datoga peoples in Tanzania and an individual from the North Plain Tribes in Montana (USA). Written informed consent was obtained from all participants. Research & ethics approvals were obtained from the MIT IRB (protocol #1612797956), but also in each sampled country prior to the start of sample collection, from the following local ethics committees: Chief Dull Knife College (Montana), protocol #FWA00020985; Comite National d’Ethique de la Recherche pour la Sante Humaine (Cameroon), protocol #2017/05/901/CE/CNERSH/SP; Nunavut Research Institute (Canada), protocol #0205217N-M; National Institute for Medical Research (Tanzania), protocol #NIMR/HQ/R.8a/Vol. IX/2657; Coordinating Ethics Committee of Helsinki and Uusimaa Hospital District (Finland), protocol #1527/2017.

Participants produced a fecal sample in a sterile container that was immediately returned to researchers in the field. Raw stool was diluted 2:10 in 25% pre-reduced (anaerobic) glycerol solution containing acid-washed glass beads, and were immediately homogenized and aliquoted into cryogenic 2ml tubes. Stool samples aliquoted in cryoprotectant were immediately flash frozen in the field at −196C, using a cryoshipper tank. Samples were then shipped to MIT for processing, culturing and storage.

Supplemental Table 1 contains metadata information about each subject enrolled in this study.

### DNA extraction, library construction and Illumina sequencing for shotgun metagenomics

We used the DNeasy PowerSoil Kit (Qiagen) with manufacturers’ protocols to extract microbial genomic DNA from stool samples. Genomic DNA libraries were constructed from 1.2ng of cleaned DNA using the Nextera XT DNA Library Preparation kit (Illumina) according to the manufacturer’s recommended protocol, with reaction volumes scaled accordingly. Prior to sequencing, libraries were pooled by collecting equal quantity of each library from batches of 94 samples. Insert sizes and concentrations of each pooled library were determined using an Agilent Bioanalyzer DNA 1000 kit (Agilent Technologies). Paired-end sequencing (2×150-bp reads) was performed using an Illumina NextSeq 500 instrument (Illumina Inc) at the Broad Institute.

### Culturing and isolation of bacterial isolates

To culture and isolate bacterial strains, we used 43 stool samples collected from 34 individuals across 9 human populations. To obtain an exhaustive representation of the diversity of human gut bacteria, human fecal samples were processed anaerobically at every step in a chamber, using gas monitors controlling physico-chemical conditions (5% Hydrogen, 20% Carbon dioxide, balanced with Nitrogen). Human fecal samples were diluted in pre-reduced PBS (with 0.1 % L-cysteine hydrochloride hydrate). Diluted samples were then plated onto pre-reduced agar plates and incubated anaerobically at 37°C for 7 to 14 days. Both general (nonselective) and selective media were used to culture diverse groups of organisms. We used 12 different media, combined with antibiotic, acid, and ethanol treatments, resulting in 19 different culturing conditions to isolate 3,632 bacterial strains from the 11 Openbiome donors. We used 6 different culturing conditions to isolate 2,556 bacterial strains from our other set of 23 individuals. See Supplementary Table 2 for culturing media used in this study. After incubation, bacteria were isolated by picking individual colonies with an inoculation loop. They were streaked onto a second pre-reduced agar plate to increase colony purity. After 2 days of incubation at 37°C, one colony was re-streaked again onto third agar plate for 2 additional days of incubation. One colony from each individual streak was then inoculated in liquid media in a 96-well culture plate. After 2 days of anaerobic incubation at 37°C, the taxonomy of the isolate was identified using 16S rRNA gene Sanger sequencing (starting at the V4 region). We first amplified the full 16S rRNA gene by PCR (27f 5’-AGAGTTTGATCMTGGCTCAG-3’ - 1492r 5’-GGTTACCTTGTTACGACTT-3’) and then generated a ~1kb long sequence by Sanger reaction (u515 5’-GTGCCAGCMGCCGCGGTAA-3’). All isolates are stored in −80°C freezers in a pre-reduced cryoprotectant glycerol buffer.

### DNA extraction, library construction and Illumina sequencing of Whole Genomes

We used the DNeasy UltraClean96 MicrobioalKit (Qiagen) and the PureLinkPro96_gDNAkit (Invitrogen) kits to extract whole genome DNA from isolate colonies, following manufacturers’ protocols. Genomic DNA libraries were constructed from 1.2ng of DNA using the Nextera XT DNA Library Preparation kit (Illumina), following the manufacturer’s protocol, with reaction volumes scaled accordingly. Prior to sequencing, we pooled on average 250 samples with equal quantities of DNA. Insert size and concentration of each pooled library were determined using an Agilent Bioanalyzer DNA 1000 kit (Agilent Technologies). Paired-end (2×150bp) reads sequencing was performed using an Illumina NextSeq 500 instrument (Illumina Inc) at the Broad Institute.

### Draft assembly and annotation of whole genome sequences

All parameters used to generate whole genome assemblies from 2×150bp paired-end data and used to perform downstream genomic analyses are embedded in the method descriptions below.

Briefly, reads were first demultiplexed using in-house scripts. We used cutadapt v1.12 ^29^ to remove barcodes and Illumina adapters (with parameters -a CTGTCTCTTAT -A CTGTCTCTTAT). We used Trimmomatic v0.36 ^30^ for the quality filtering of data (with parameters PE -phred33 LEADING:3 TRAILING:3 SLIDINGWINDOW:5:20 MINLEN:50). Reads were assembled de novo into contigs using SPAdes v.3.9.1 ^31^ (with parameter--careful). To iteratively improve genome assemblies, we used SSPACE v3.0 ^32^ and GapFiller v1-10 ^33^ to scaffold contigs and to fill sequence gaps (with default parameters). Scaffolds smaller than 1kb were removed from genome assemblies. We aligned all reads back to the assembly to compute genome coverage using BBmap v37.68 (https://jgi.doe.gov/data-and-tools/bbtools/) and the covstats option (with default parameters). The final assemblies were annotated using Prokka v1.12 ^34^ (with default parameters).

### Assessing assembly quality

We measured genome assembly statistics using CheckM v1.0.7 ^35^ (with parameters lineage_wf --tab_table -x fna Prokka_annotations/). Next, we used the Strucchange R package to remove contaminant contigs. Contaminations are often characterized by small contigs with extreme coverage. Thus, we detected breakpoints in the distribution of sorted coverages across contigs ^36^ (with cov defined as a sorted vector of contig converages, the function breakpoints(log(cov)~seq(1,length(cov))) was used to calculate the breakpoints). If multiple jumps in coverage data are detected, the contig with the highest coverage is selected as the breakpoint. Then, all contigs with higher or equal coverage to the breakpoint contig are excluded from the assembly file. Finally, we conserved all assemblies that had genome completeness higher than 90%. All summary and quality statistics can be found in Supplementary Table 3. The median assembly completeness of all 6,188 genomes is 99.41%, the median contamination is 0.3%, the median coverage is 145kb, and the median coverage is 126X.

### Clustering genomes into species

We used whole genomic information to group genomes into species clusters. We used an open-reference approach and computed all-against-all genomic distances using Mash ^37^ (with default parameters). A Mash distance lower than 0.05 is equivalent to using an Average Nucleotide Identity higher than 95 %, which is a standard threshold for delineating species ^38^. We used an unsupervised hierarchical clustering approach to group genomes that had Mash distances <= 0.05 into taxonomic units using the bClust function from the micropan R package ^39^. We then measured the genetic distance between the representative genome of each species cluster (defined as the genome with the highest N50) and 79,226 non-contaminated complete and draft genomes downloaded from the NCBI FTP repository (ftp://ftp.ncbi.nlm.nih.gov/genomes/) on March 27th, 2017. Clusters with a Mash distance to NCBI genomes lower than 0.05 was assigned the taxonomy of the closest reference genome (we manually curated Mash results to assign a taxonomy to each cluster when NCBI taxonomies were incomplete or incorrect). All genome taxonomies are compiled in Supplementary Table 3.

### Detection of HGTs

We looked for gene transfers that occurred between genomes of different bacterial species. We used Blast (blastn, v2.6.0) ^40^ to systematically detect blocks of DNA that are shared by two genomes. We retained blast hits with 100% similarity and that are larger than 500bp. To increase the likelihood of looking at transfer events that occurred on timescales compatible with human lifetime, we focused many of our analyses (Figures 1B-E) on transferred blocks that are larger than 10kb. We removed blast hits that involve contigs with k-mer assembly coverage lower than 3.

### Calculating HGT frequencies

To measure the frequency of HGT between two species, we only considered the fraction of genome pairs that share at least one HGT. To avoid inflating estimations of HGT frequency, we did not consider the absolute number of distinct blast hits between two genomes, as poor assembly or genomic processes, such as transposition, might result in splitting a single large mobile element into many smaller apparent HGT events.

### Abundance of species and genomes

Because bacterial species abundance can vary across people, we measured average species abundances within each individual host. For species with more than 5 isolate genomes per individual, we randomly selected 5 genomes to compute the average abundance. For species with less than 5 isolate per individual, we used all isolates to calculate the average abundance. We mapped metagenomic data generated from the same individual host against each isolate genome, and used the per base coverage K, the average read length L, the size of each genome S and the total number of reads T in the shotgun data to calculate the relative abundance A of each genome in the metagenome with A = (K*S/L) / T. We used a threshold of 0.01 to define lowly and highly abundant bacteria.

### Assigning Gram stain to bacterial species

We used Gram staining data from reference microbiology databases (ATCC (http://www.lgcstandards-atcc.org/en.aspx), DSMZ (https://www.dsmz.de/) & the Microbe Directory database (https://microbe.directory)) and from publications characterizing the phenotype of bacterial isolates to assign a consensus Gram stain to each of our bacterial species. Species with contradictory Gram staining information or with unknown taxonomy were excluded from the analysis of the correlation between HGT frequency and cell wall architecture. Our data recapitulate what we know from the literature ^41,42^: Bacteroidetes are Gram-; Bifidobacterium are Gram+; Firmicutes are Gram+, to the exception of Negativicutes species, which are known diderm bacteria, and of a few other species; Fusobacterium are Gram-; Akkermansia are Gram-; Proteobacteria are Gram-.

### Annotating transferred genes

Functional annotation followed the basic approach described previously ^24^. Briefly, CDS were assigned to mobile gene contigs at least 500 bp in length using Prodigal ^24,43^ in metagenome mode to capture gene fragments. The resultant CDS were dereplicated and clustered at 90% nucleotide identity using vsearch ^44^. These gene centroids were used for subsequent functional annotation steps. Both eggNOG-mapper ^45^ and InterProScan ^46^ were used to assign putative function predictions to gene centroids. For additional classification of antibiotic resistance genes and carbohydrate active enzymes, hmmer3 ^47^ was used with the Resfam ^48^ and dbCAN ^49^ hmm databases with a cutoff e-value of 1e-5 and score of 22. Text mining with a set of regular functional annotations that we previously used ^24^ was employed to determine the assignment of genes into the following categories: phage, plasmid, transposons, and antibiotic resistance.

### Statistical analyses

Statistical analyses were performed in R. We compared HGT frequencies within individuals vs. between individuals for the same species pairs, by comparing the observed total number of genome comparisons with at least 1 HGT within people to its expected value, and calculated p-values from the Poisson distribution (ppois R function). The expected total number of genome comparisons with at least 1 HGT within people was calculated based on HGT frequencies found between people. The same approach was used to compare HGT frequencies of the same species pairs found in the US population vs. the non-industrialized populations. The individual contributions of phylogeny, species abundance and cell wall architecture to within-person HGT frequency were measured using a linear mixed effects model, assuming an intercept that is different for each host population. We used the lmerTest R package ^50^ (lmer function), which provides *p-values* for linear mixed effects model fits. Profiles of COG functional categories were compared using a chi-square Goodness-of-fit test (chisq.test function). HGT counts of phage, plasmid, transposon, antibiotic resistance and CAZyme genes were compared between industrialized and non-industrialized host populations using two-proportions Z-tests (prop.test function), and a Bonferroni correction for multiple tests (p.adjust function).

