## Supplemental Material for "Industrialization is associated with elevated rates of horizontal gene transfer in the human microbiome"

### Supplementary Information

#### High within-people HGT frequency mainly reflects transfers that occurred within the host of origin.

The criteria that we used to detect HGTs restrict the timescale of these events to a time window compatible with a human lifespan. We cannot exclude the possibility that some of these transfers occurred one or two generations ago in the host's parents or grandparents. However, it is unlikely that such scenarios would drive the difference of HGT frequency that we observe across all bacterial species pairs that we sampled within and between people. For such ancient transfers to be seen in contemporary adult microbiomes, faithful co-transmission of pairs of strains involved in a given HGT from one generation to another, followed by persistent colonization until adulthood, must occur.

It was recently shown that family members or individuals living in the same household tend to share more strains together than with unrelated individuals, or than with people living in different houses [1–3]. Yet, the percentage of individual strains that are shared between two people, resulting either from horizontal (e.g. within a household) or vertical transmission (e.g. at birth), is relatively small compared to the whole diversity of bacterial strains colonizing a gut microbiome [1–3]. For example, within a cohort of 25 healthy women, the rate of mother-to-infant strain transmission was estimated to be about 16% [1]. The gut (mother) to gut (infant) transmission rate is actually even lower than 16%, as this estimation accounts for transmission of strains from multiple body sites in the mother [1].

As a result, the percentage of pairs of strains that are shared by any two people must be much smaller (*e.g.*, 0.16 * 0.16 = 1.3%). The number of strain pairs that also co-exist into adulthood is likely even smaller [4–6]. By comparison, the within-individual rate of HGT for species separated by a distance of 0.2 amino acid subs/site is nearly 10%.

Moreover, in a recent study, we found that the residence time of *Bacteroides fragilis* strains is around ~1 year for the majority of strains in an adult gut microbiome and about 10 years for the longest resident, showing that strain composition can rapidly change.

Because the co-occurrence of identical strains in different people is so rare, and that strain turnover is frequent from birth to adulthood, it is highly probable that the higher frequency of within person HGT compared to between person HGT is explained by gene transfers that occurred within the gut of sampled individuals.

### Supplementary Figures

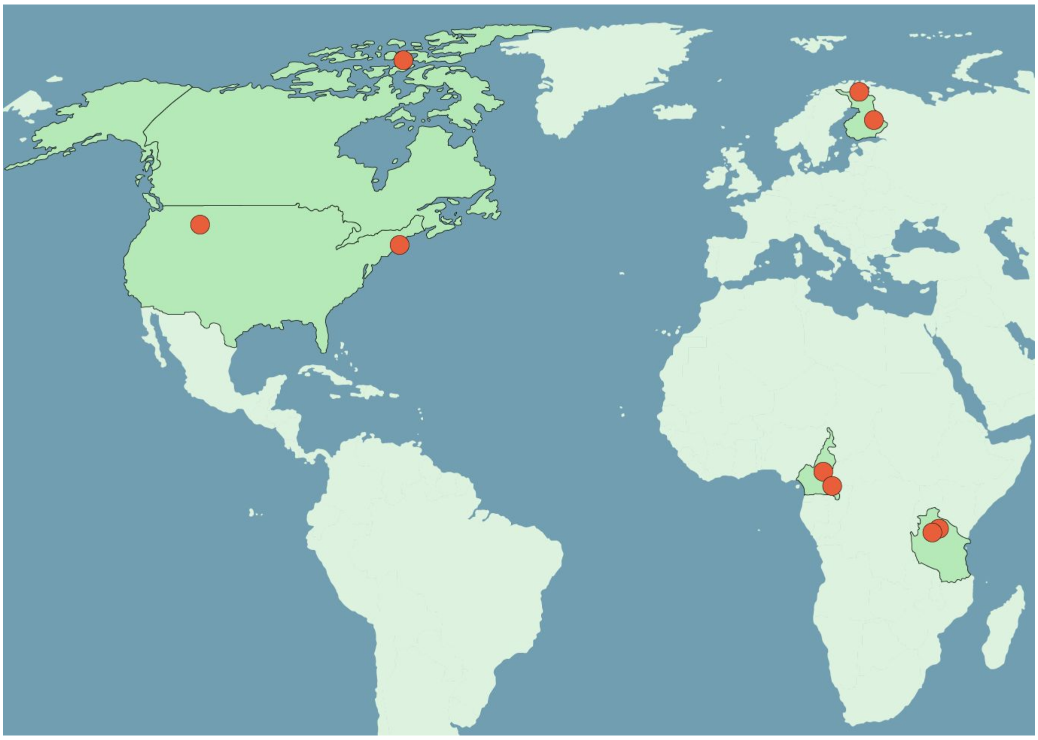

**Supplementary Figure 1 - Sampled locations and populations**

Samples were collected among 9 communities living in the USA, Canada, Finland, Cameroon and Tanzania. Red dots show the geographic locations of sampling sites. We recruited individuals with different lifestyles and diets: industrialized with western diets in the Boston area (USA), in the Northern Plain Tribes in Montana (USA), and in eastern (Finnish) & arctic (Sami) Finland ; Non-industrialized with semi-traditional diets in Canadian arctic (Inuits); non-industrialized with traditional diet in Cameroon (Beti, agriculturalists and Baka, hunter-gatherers/farmers); non-industrialized with traditional diets in Tanzania (Datoga, pastoralists and Hadza, hunter-gatherers). See Supplementary Table 1 for more detailed information on each individual/population.

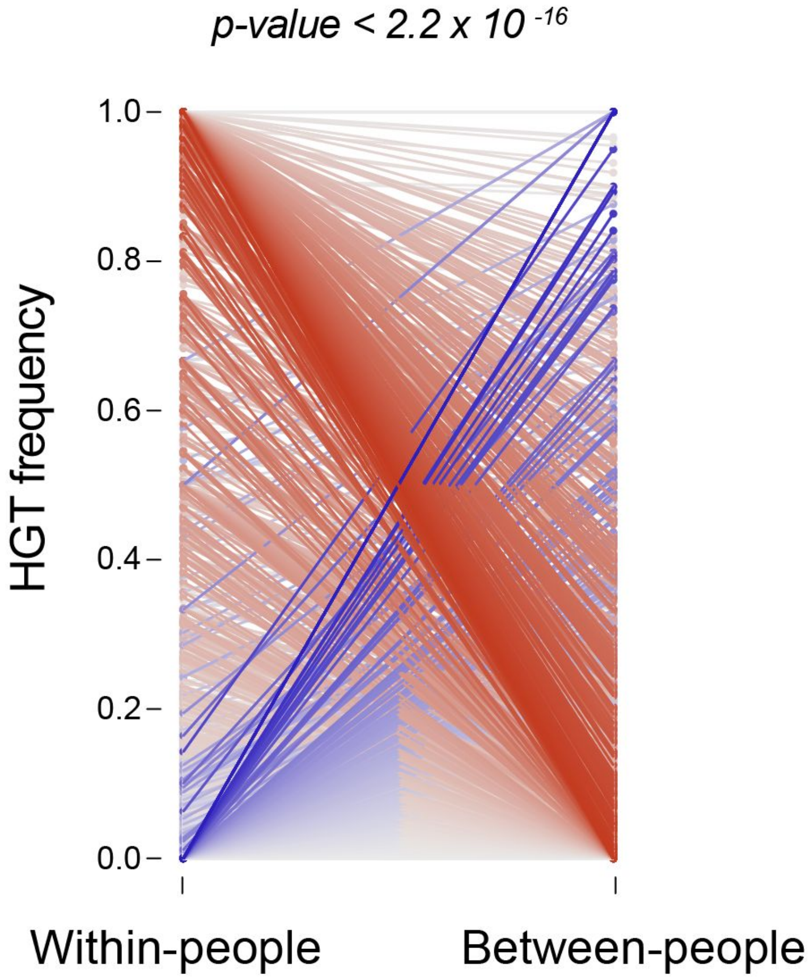


**Supplementary Figure 2 - Extensive gene transfers in the gut microbiome of each individual, using mobile elements larger than 500bp.**

These results replicate findings of Figure 1A and B derived from the analysis of all 134,958 mobile elements larger than 10kb. HGT frequencies within and between people are shown for all bacterial species pairs and are represented by solid lines. HGT frequencies were computed from all 5,126,962 mobile elements that are larger than 500bp. Differences in HGT frequency are colored along a gradient from grey (no difference) to red (within-people HGT frequency is higher than between-people) or from grey to blue (between-people HGT frequency is higher than within-people), darker colors representing higher differences. The HGT frequency of bacterial species pairs found within people were compared to their expected HGT frequency based on the HGT frequency of the same species pairs found between people. P-values were calculated using a Poisson distribution. Observed and expected HGT frequencies were calculated using the total number of genome comparisons with at least 1 HGT (see Methods).

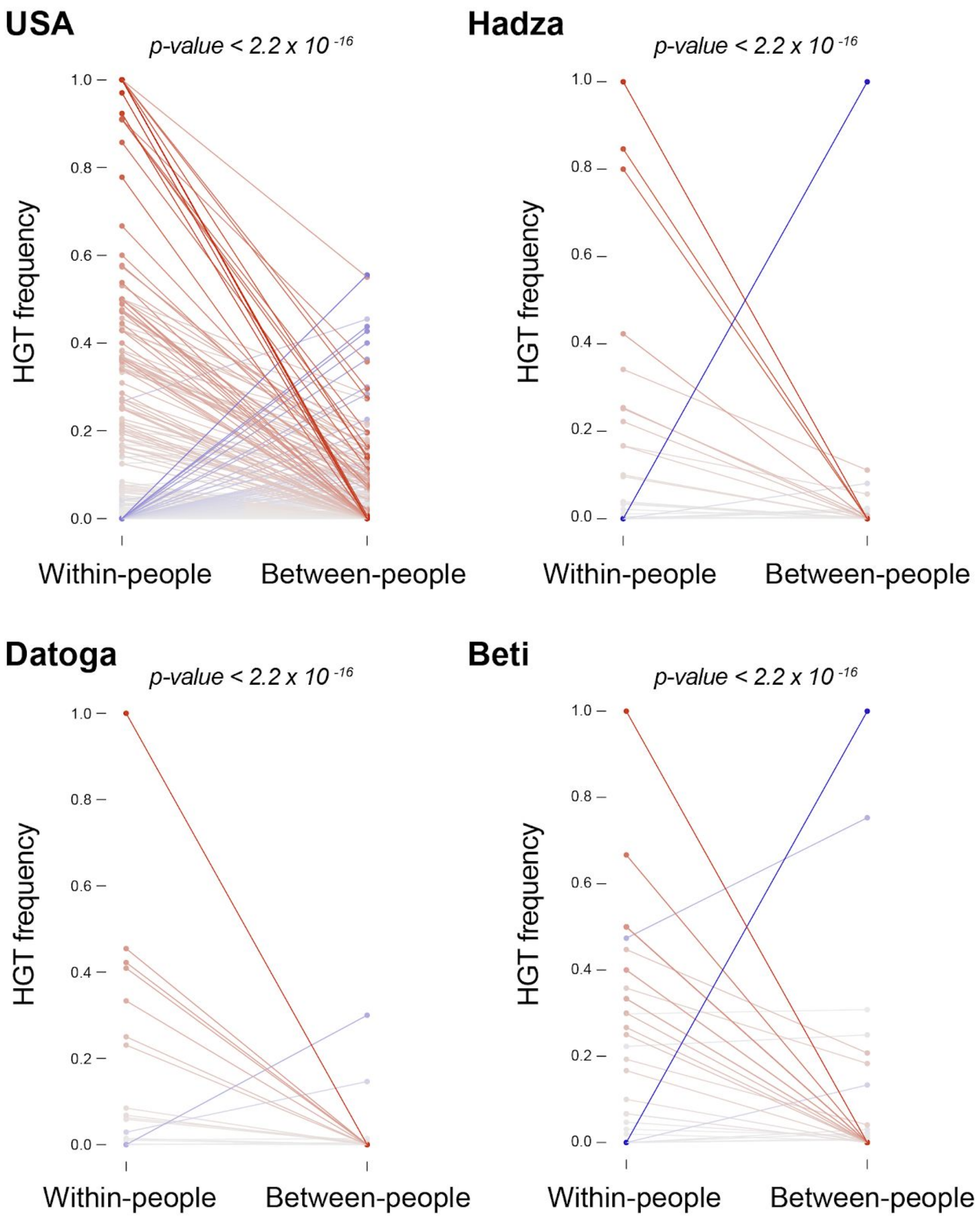


**Supplementary Figure 3 - Gene transfers in the gut microbiome of each individual within each sampled population.**

Increased HGT frequency within individuals is also found when comparing within- to between-people HGT frequencies computed within each of our populations in which we had sufficient sampling size. As in Figure 1B, species pairs are represented by solid lines. HGT frequencies were computed from all mobile elements that are larger than 10kb. Differences in HGT frequency are colored along a gradient from grey (no difference) to red (within-people HGT frequency is higher than between-people) or from grey to blue (between-people HGT frequency is higher than within-people), darker colors representing higher differences. The HGT frequency of bacterial species pairs found within people were compared to their expected HGT frequency, based on the HGT frequency of the same species pairs found between people. P-values were calculated using a Poisson distribution.

Observed and expected HGT frequencies were calculated using the total number of genome comparisons with at least 1 HGT (see Methods).

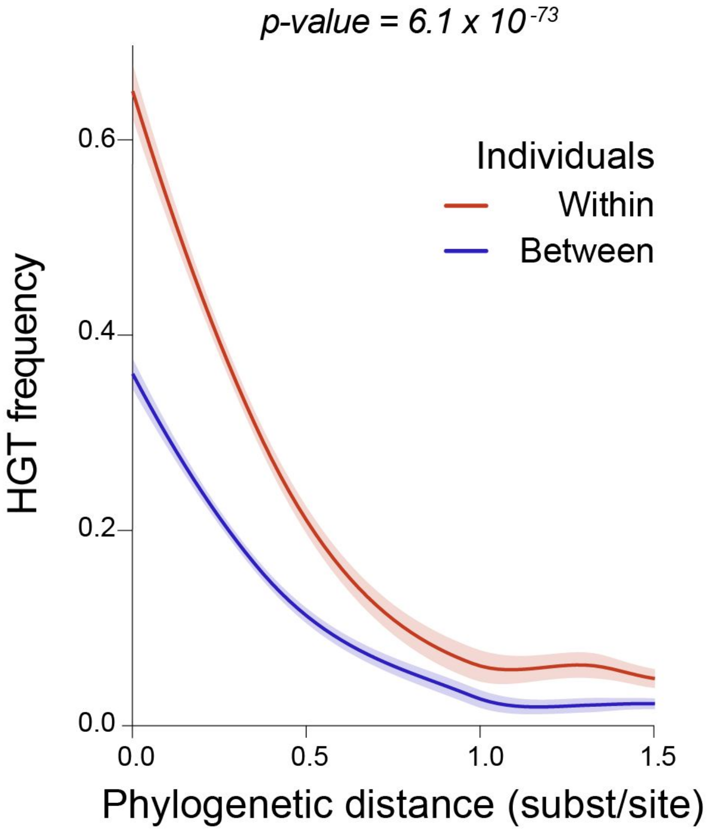


**Supplementary Figure 4 - Mobile elements larger than 500bp show that within-people HGT occurs at higher rates than between-people across all phylogenetic distances.**
All 5,126,962 mobile elements that are larger than 500bp were used to calculate the HGT frequency for all species pairs sampled both within people and between people. The HGT frequency is plotted against the phylogenetic distance between bacterial species. Within people HGT frequency is higher than between people across all phylogenetic distance bins. Phylogenetic distances were derived from the phylogenomic tree in Figure 1A.

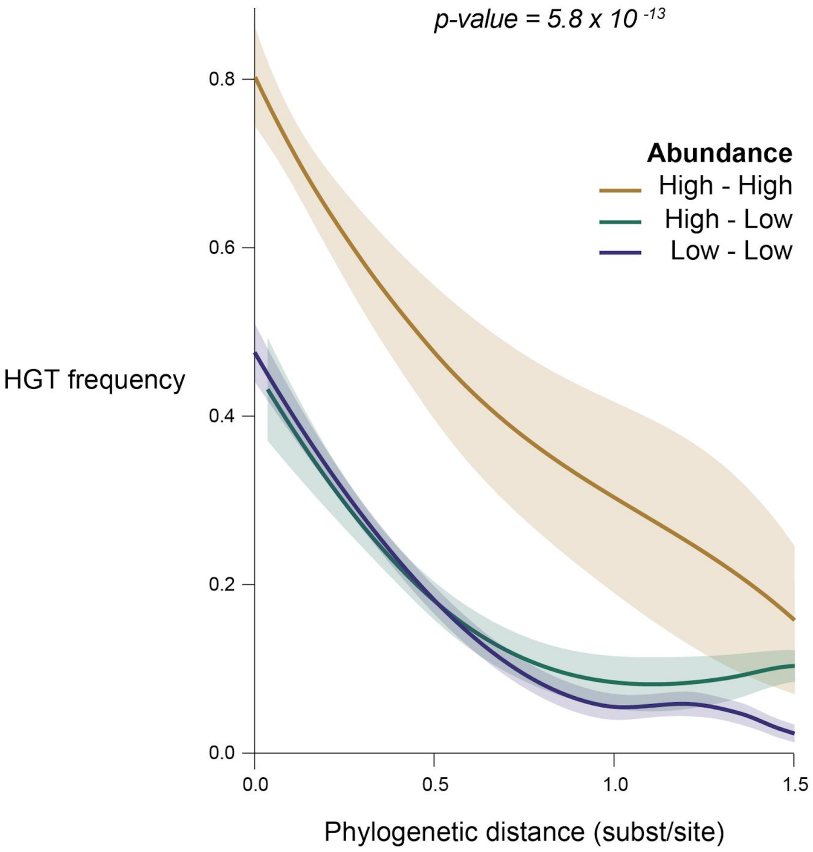


**Supplementary Figure 5 - Mobile elements larger than 500bp show that bacterial abundance is driving HGT within people.**

All 5,126,962 mobile elements that are larger than 500bp were used to calculate the HGT frequency for all species pairs sampled within people. HGT frequency is plotted across species abundance bins. Bacterial abundance was measured for all species in each individual, by mapping metagenomic reads against individual genomes (see Methods). We used a threshold of 0.01 to define highly and lowly abundant bacteria. HGT frequencies are plotted against the phylogenetic distance between bacterial species. Phylogenetic distances were derived from the phylogenomic tree in Figure 1A. HGT frequency is plotted using loess regression, with confidence intervals being calculated from the standard errors. The individual contribution of abundance, independent of phylogeny and cell wall architecture, was measured using a Linear Mixed Effects model. The p-value associated to the specific contribution of abundance is shown above the plot.

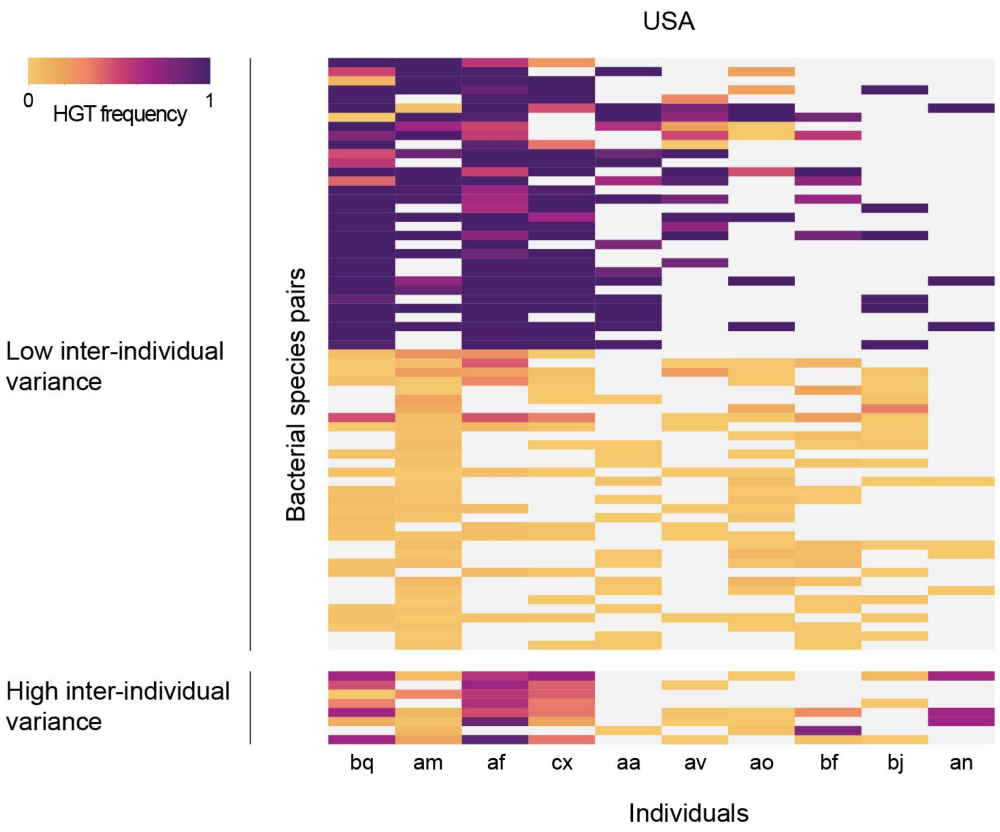


**Supplementary Figure 6 - The same bacteria exchange genes in the gut microbiome of different individuals.**

We compared within-individual HGT frequencies for bacterial species pairs shared by a minimum of 4 individuals, using data from our USA cohort (other populations were not considered because of limited numbers of species pairs that are shared across multiple individuals). HGT frequencies are represented in a heatmap, with individuals in columns and species pairs in rows. The color gradient represents HGT frequency, from yellow (no transfer observed) to purple (maximum frequency). To measure homogeneity of transfer rates for each species pair across individuals, we used a randomization test. We computed all standard deviations of HGT frequency, and compared the observed average standard deviation to a null distribution of average standard deviations derived from randomly reshuffling HGT frequencies in the data (*p-value* $< 0.001$).

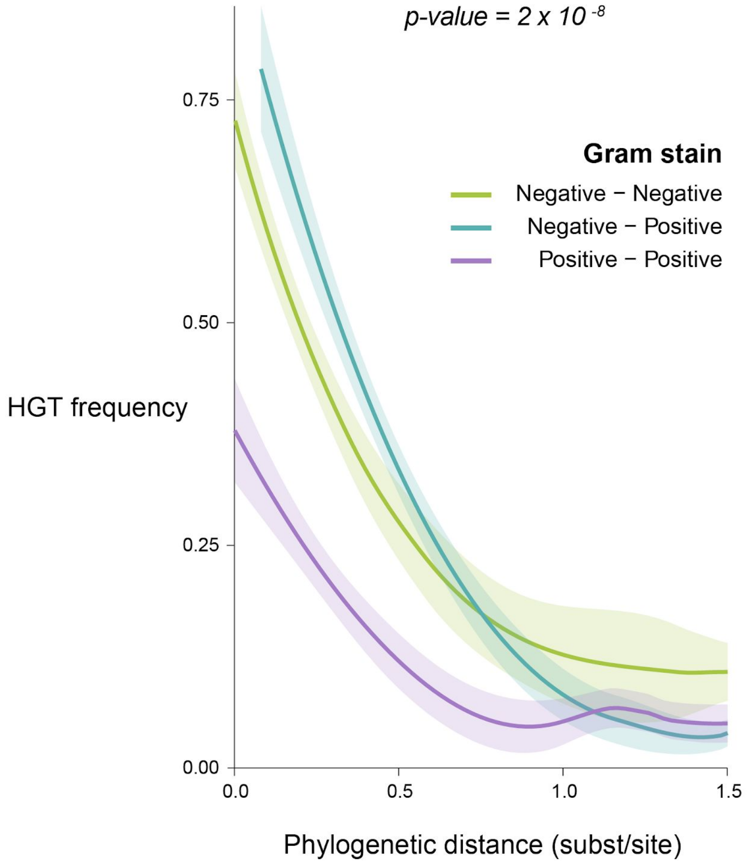


**Supplementary Figure 7 - Mobile elements larger than 500bp show that bacterial cell wall architecture is driving HGT within people.**

All 5,126,962 mobile elements that are larger than 500bp were used to calculate the HGT frequency for all species pairs sampled within people. HGT frequency is plotted across types of cell wall architecture. We used Gram staining as a proxy to call for monoderm or diderm bacteria. (see Methods). HGT frequencies are plotted against the phylogenetic distance between bacterial species. Phylogenetic distances were derived from the phylogenomic tree in Figure 1A. HGT frequency is plotted using loess regression, with confidence intervals being calculated from the standard errors. The individual contribution of cell wall architecture, independent of phylogeny and abundance, was measured using a Linear Mixed Effects model. The p-value associated to the specific contribution of cell wall architecture is shown above the plot.
